# Dynamic changes in uterine NK cell subset frequency and function over the menstrual cycle and pregnancy

**DOI:** 10.1101/2022.02.21.481277

**Authors:** Emily M Whettlock, Ee Von Woon, Antonia O Cuff, Brendan Browne, Mark R Johnson, Victoria Male

## Abstract

Uterine Natural Killer cells (uNK) play an important role in promoting successful pregnancy by regulating trophoblast invasion and spiral artery remodelling in the first trimester. Recently, single cell RNA sequencing (scRNAseq) on first trimester decidua showed that uNKs can be divided into three subsets, which may have different roles in pregnancy. Here we present an integration of previously published scRNAseq datasets, together with novel flow cytometry data to interrogate the frequency, phenotype and function of uNK1-3 in seven stages of the reproductive cycle (menstrual, proliferative, secretory phases; first, second, third trimester; and postpartum). We found that uNK1 and −2 peak in the first trimester, but by the third trimester the majority of uNK are uNK3. All three subsets are most able to degranulate and produce cytokines during the secretory phase of the menstrual cycle and express KIR2D molecules, which allow them to interact with HLA-C expressed by placental extravillous trophoblast cells, at the highest frequency during the first trimester. Taken together, our findings suggest that uNK are particularly active and able to interact with placental cells at the time of implantation, and that uNK1 and 2 may be particularly involved in these processes. Our findings are the first to establish how uNK frequency and function changes dynamically across the healthy reproductive cycle. This serves as a platform from which the relationship between uNK function and impaired implantation and placentation can be investigated. This will have important implications for the study of subfertility, recurrent miscarriage, pre-eclampsia and pre-term labour.

## Introduction

Uterine Natural Killer cells (uNK) are NK-like cells that are found in the lining of the uterus (called “decidua” in pregnancy and “endometrium” outside of pregnancy). They are distinct from peripheral blood NK (pNK) in both phenotype and function. Unlike pNK, uNK are only weakly cytotoxic and instead produce factors that are pro-angiogenic and that attract fetally-derived placental cells called extravillous trophoblast (EVT) (1–6). uNK are most prominent in the first trimester of pregnancy, at which time they account for 70-80% of immune cells in the decidua (7). Their prominence at the time of implantation, and production of factors that are predicted to promote trophoblast invasion and spiral artery remodeling, indicates they are likely to have a role in placental implantation.

Further evidence for a role of uNK in implantation comes from their expression of high levels of killer-cell immunoglobulin-like receptors (KIRs), CD94/NKG2 (8, 9), and LILRB1 (9), which allows them to recognize the human leukocyte antigens (HLAs) expressed by EVT: HLA-C, HLA-E and HLA-G respectively (10–12). Immunogenetic studies demonstrate that combinations of HLA and KIR that lead to lower activation are associated with disorders of insufficient implantation such as pre-eclampsia, fetal growth restriction and recurrent miscarriage, suggesting that uNK activation via KIR is important for implantation (13–20). The increased expression of KIRs by uNK around the time of implantation provides additional support for this idea (21).

Until recently, it was thought that uNK formed a single population, but single cell RNA sequencing (scRNAseq) has now demonstrated three subpopulations of uNKs in first trimester decidua (22). These were originally called decidual NK (dNK) −1, −2 and −3 but they have now also been found in non-pregnant endometrium (23). Here, we call these subsets “uNK1”, “uNK2”, and “uNK3” in recognition of the finding that they are not confined to the decidua. uNK1 express higher levels of KIRs and LILRB1, indicating they may be specialized to communicate with EVT (22, 24). uNK2 and uNK3 produced more cytokines upon stimulation, indicating their role may be immune defense (24). However, several questions remain open. Are these three subpopulations still present at the end of pregnancy? Do the subpopulations change in prominence and/or activity over the reproductive cycle? The answers to these questions could elucidate which subpopulations are important in implantation, parturition, and immune protection throughout the reproductive cycle.

Here, we show that the proportions of the uNK subpopulations remain stable through the menstrual cycle, but all three are more active and express higher levels of KIR around the time of implantation. uNK1 are more prominent in the first trimester of pregnancy, potentially indicating a requirement for this subset in the mediation of implantation whereas uNK3 are the most prominent at the end of pregnancy. Overall, we outline how the three uNK subpopulations change in proportion phenotype and function throughout the reproductive cycle.

## Materials and Methods

### Primary tissue

Collection of human tissue was approved by London - Chelsea Research Ethics Committee (study numbers: 10/H0801/45 and 11/LO/0971).

29 endometrial samples were taken by Pipelle biopsy before insertion of intrauterine device for contraception. Samples taken on a day of bleeding were assigned as menstrual phase. Other samples were categorized to proliferative or secretory phase by date of last menstrual period and serum progesterone level. Samples obtained before Day 14 were assigned as proliferative phase and after day 14 as secretory phase. This was confirmed retrospectively by serum progesterone level, according to previously published reference range (25). Samples from postpartum participants were assigned by number of days from delivery of the baby up until 16 weeks post-partum.

10 decidual samples were taken from participants undergoing surgical management of elective termination of pregnancy between 6 to 13 weeks of pregnancy and 10 from participants undergoing elective caesarean sections, over 37 weeks of pregnancy and not in labour. For labouring data, a following 8 samples were taken from participants in the early stages of labour (1-3cm cervical dilation and regular contractions) who had a caesarean section, and 5 samples were taken from participants after a vaginal birth. Matched peripheral venous blood was obtained from all patients at time of obtaining endometrial or decidual samples. Patient characteristics are summarized in Suppl tables 1 and 2.

Lymphocytes were extracted from peripheral blood by layering onto Histopaque (Sigma Aldrich), spinning down (700 *xg*, 20 minutes, 21°C) and retrieving the interface which was washed twice with Dulbecco’s Phosphate-Buffered Saline (Life Tech) (500 *xg*, 10 minutes, 4°C). Briefly, endometrial tissue was passed through 100μm cell strainer, pelleted (700 *xg*, 10 minutes, 4°C), resuspended in Dulbecco’s Phosphate-Buffered Saline supplemented by 10% Fetal Calf Serum (Sigma Aldrich), passed through 70μm strainer, and layered on Histopaque as above.

For first trimester samples, decidua compacta was extracted from products of conception and stirred for 20 minutes to remove blood before mincing with scalpel followed by GentleMACS dissociation (Miltenyi). Minced tissue was passed through 75 μm sieve, pelleted (500 *xg*, 10 minutes, 4°C) and resuspended in PBS/1% FCS before passing through 100 μm strainer. Filtrate was layered on Histopaque as above.

For third trimester decidua basalis (DB) samples, small sections were cut from the maternal side of the placenta and washed using a magnetic stirrer in Mg^2+^ and Ca^2+^ free PBS (Gibco) for 20 minutes. Blood clots, vessels and placental tissue were physically removed and cleaned decidual tissue was placed in new Mg^2+^ and Ca^2+^ free PBS. The tissue was spun (400 *xg*, 5 mins 21°C) and PBS removed. The tissue was resuspended in Accutase (Invitrogen), mechanically digested in C tubes using a GentleMACs dissociater and placed in a 37°C shaking water bath for 45 minutes. Minced tissue was passed through a 70μm strainer, resuspended in PBS/1%FCS/2mM EDTA. Filtrate was layered on Histopaque as above. For the third trimester decidua parietalis (DP) samples, 10cm x 10cm sections of the fetal membrane were dissected and the decidua removed using a cell scraper (Starsted). The tissue underwent enzymatic and mechanical digestion as described in the decidua basalis protocol.

Extracted lymphocytes were counted by light microscopy (Leica) with a haemocytometer. A total of 0.2 X10^6^ to 1 X10^6^ cells per condition were allocated for phenotype and functional assessment.

### Stimulation with PMA/ionomycin

21 endometrial, all first trimester and all third trimester samples were used for functional assessment. Endometrial lymphocytes were stimulated immediately after isolation, and decidual lymphocytes were stimulated after 12 to 20 hours of rest at 37°C. Optimization experiments showed no difference between cells stimulated fresh and after rest (Supp Fig 1).

For functional assessment, cells were suspended in RPMI enriched with antibiotics, EDTA and sodium pyruvate and divided into unstimulated and stimulated wells. Anti-CD107a BV605 (100 μl/ml), Brefeldin (10μg/ml) and Monensin (2μM/ml) were added to all wells and Phorbol 12-myristate 13-acetate (PMA) (50 ng/ml) and ionomycin (1μg/ml) into the stimulated wells only. Cells were incubated for 4 hours at 37°C then stained with antibodies. For third trimester samples, cells were incubated for 6 hours with anti-CD107a, PMA and ionomycin, with Brefeldin and Monensin added 2 hours into the incubation.

### Single cell RNA-seq data analysis

For the scRNAseq data, R(26) was used for the majority of the analysis. This included use of the package Seurat (27), designed for analysis of scRNAseq data, and various data manipulation and visualisation packages (28–35).

A scRNA seq dataset from the non-pregnant uterus (available at www.reproductivecellatlas.org/) was converted from a Python format into a R Seurat object. The object was subset to cells that had been classified as “Lymphoid” or “Myeloid” under “Cell.type”, in the metadata.

A scRNAseq dataset from the first trimester uterus (available at Array Express E-MTAB-6701) was converted from .txt into a R seurat object. The object was subset to cells that had been designated an immune cell type, e.g. “dNK1” under “Annotation”, in the metadata. Cells originating from placenta or blood were removed, so only decidual cells remained. Data from one donor was removed from the analysis due to their NK cells clustering independently of all other NK cells.

The scRNAseq dataset (dbGaP phs001886.v1.p1, reanalyzed with permission of the NIH, project ID 26528) contained samples from one second trimester accreta sample and 9 third trimester participants. Both datasets were filtered, aligned and quantified using Cell Ranger software (version 5.0.1, 10x Genomics). h19 was used as a human genome reference. Downstream analyses were performed using the R package Seurat(27). Cells with fewer than 200 genes and genes that were expressed in less than 10 cells were removed. Furthermore, cells where the gene content was greater than 10% mitochondrial genes were removed. Clusters were identified using “FindClusters” algorithm. The “FindAllMarkers” algorithm was used to identify the immune clusters, and subset the object to immune cells. Cells from placental tissue were removed. TIL and PTL samples were not included in the analysis looking at dNK across the reproductive cycle. PTL samples were not included in the term labouring analysis.

The four datasets were integrated based on a previously published workflow (36). Clusters that appeared to be non-immune cells were removed and the remaining cells were reanalysed using the same workflow. The algorithm “FindConservedMarkers” was used to identify the clusters dNK1, dNK2 and dNK3. This was confirmed by the metadata column “annotation” from the first trimester dataset. For visualisation of clusters and gene expression across the reproductive cycle, each dataset was downsampled so the total number of cells displayed was equal in each dataset.

### Flow cytometry

The following anti-human antibodies were used: Anti-CD56 Brilliant Violet (BV) 650 (clone NCAM 16.2, BD Bioscience), anti-CD39 BV421 (clone A1, Biolegend), anti-CD3 BV711 (clone SK7, Biolegend), anti-CD103 BV785 (clone Ber-ACT8, Biolegend), anti-CD16 Alexa Fluor(AF)700 (clone 3G8, Biolegend), anti-CD9 phycoerythrin(PE)/Dazzle 594 (clone HI9a, Biolegend), anti-CD49a PE/Cy7 (clone TS2/7, Biolegend), anti-CD45 allophycocyanin (APC) (clone HI30, Biolegend), anti-CD94 PE (Clone HP-3D9, BD Bioscience), anti-CD158a/h (KIR2DL1/DS1) VioBright 515 (clone REA1010, Miltenyi Biotec), anti-CD158b (KIR2DL2/DL3) APC vio 770 (clone REA 1006, Miltenyi Biotec), CD85j (ILT2 or CD94) Peridinin chlorophyll protein (PerCP)-eFluor 710 (clone HP-F1, Thermo Fisher Scientific) and anti-CD107a BV605 (clone H4A3, Biolegend) for surface antigens, and anti-IL-8 PE (clone G265-8, BD Bioscience), anti-IFN-Ꝩ APCvio770 (clone REA600, Miltenyi Biotec), anti GM-CSF PERCP/Cyanine 5.5 (clone BVD2-21C11, Biolegend), anti-TNFα FITC (clone MAb11, Biolegend) for intracellular staining. Cells were first incubated with fixable viability dye (Live/Dead Fixable Aqua Dead Cell stain kit, LifeTech) (15 minutes, 4°C) followed by incubation with surface antibodies (15 minutes, 4°C). For intracellular staining, human FoxP3 buffer (BD Bioscience) was used according to manufacturer’s instructions before staining with intracellular antibodies (30 minutes, 4°C). For third trimester samples, fixable viability dye was included with the surface staining antibodies (20 minutes, RT) and intracellular staining used the Cytofix/Cytoperm kit (BD Biosciences) according to manufacturer’s instructions. Excess antibodies were washed off (5 minutes, 500 *xg*, 4°C) between each incubation and twice after final incubation with intracellular antibodies.

### Statistical analysis

Data were acquired on an BD Fortessa and analysed using FlowJo (Tree Star, Ashalnd, OR). Application settings were used to ensure reproducible results. Statistical analysis were performed using PRISM (GraphPad Software Inc.). Data were assessed for normality using Shapiro-Wilk tests to determine whether a parametric or a non-parametric statistical test was appropriate. The appropriate statistical test was used to compare subsets as specified in figure legends. p<0.05 was considered significant.

## Results

We examined tissues and collected data at seven stages of the reproductive cycle. In the menstrual cycle there are three stages: menstrual (when the lining of the uterus is shed), proliferative (prior to ovulation) and secretory (after ovulation). Pregnancy is divided into three trimesters: first (1-12 weeks), second (12-28 weeks), third (28 – 40 weeks). We also examined postpartum samples (up to 16 weeks post-delivery). For the scRNAseq data, 5 stages were examined: proliferative, secretory, first trimester, second trimester and third trimester. For flow cytometry, 6 stages were examined: menstrual, proliferative, secretory, first trimester, third trimester and postpartum.

During the menstrual cycle stages the uterine tissue we examined is known as the endometrium. During pregnancy, this tissue undergoes a process called decidualization and results in three tissues known as the decidua basalis (which lines the maternal side of the placenta), decidua parietalis (which lines the rest of the uterus) and decidua capsularis (which lines the embryo on the luminal side). When taking samples from first trimester tissue, it not possible to differentiate between the different decidual tissues. During second trimester the decidua capsularis fuses with the decidua parietalis. When taking samples from third trimester tissue, it is possible to get distinct samples from the decidua basalis and the decidua parietalis.

### uNK1, −2 and −3 are present throughout the human reproductive cycle and vary in frequency

scRNAseq analysis has previously identified that three subpopulations of uNK, uNK1, −2 and −3 present in first trimester decidua (22) and non-pregnant endometrium (23). Previous analysis of scRNAseq data from third trimester decidua identified only a single cluster within the uNK population (37), and our reanalysis of the third trimester dataset alone confirmed this. However, when the third trimester data was integrated with data from the non-pregnant uterus, first and second trimester, the third trimester dNK cells did form three clusters (Fig. 1a, b).

**Figure 1.**
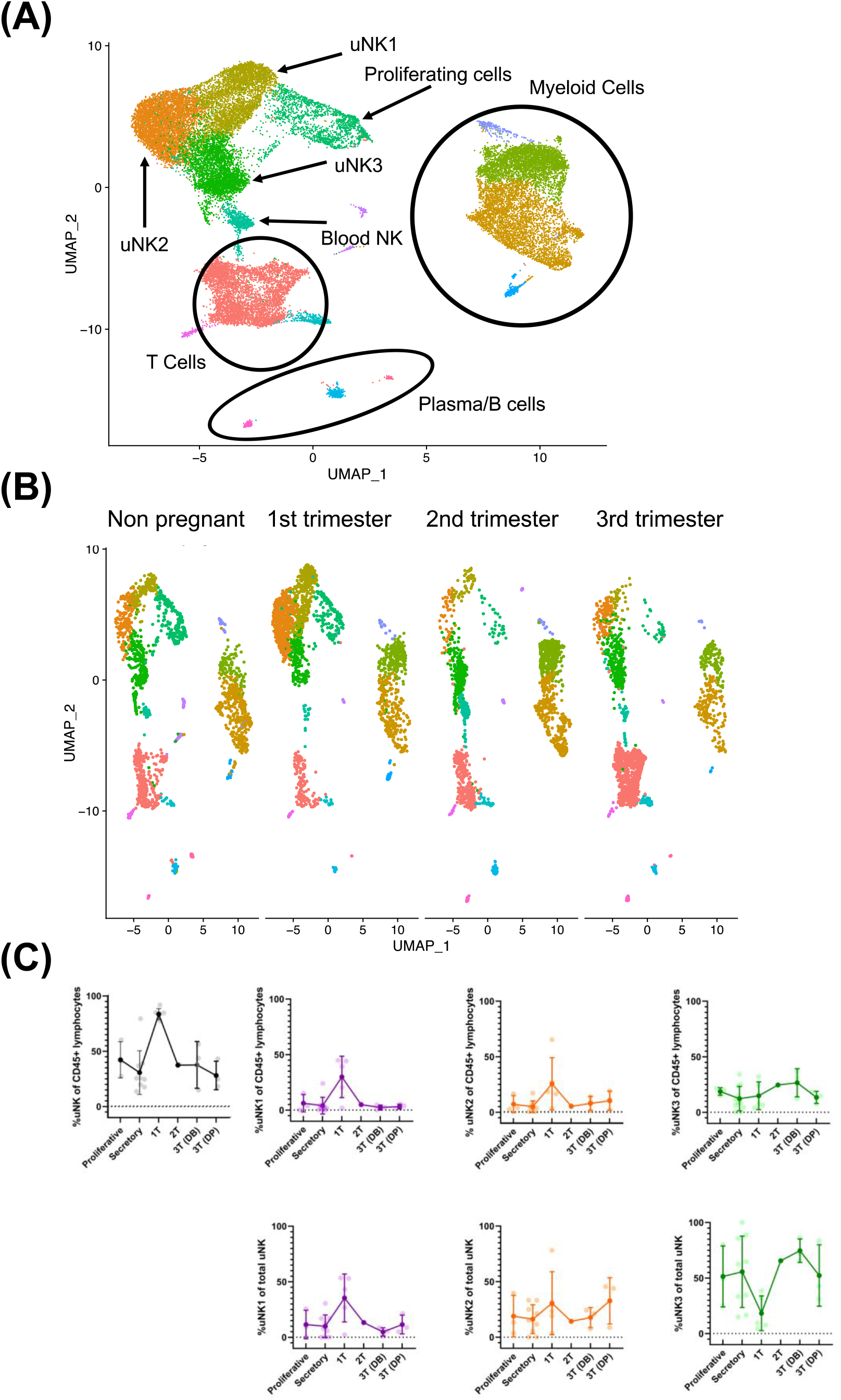
uNK1, −2 and −3 are present throughout the human reproductive cycle, by scRNAseq. **(A**) Integrated immune cells from non-pregnant endometrium, and first, second and third trimester decidua, visualised by UMAP. Colours are indicative of clusters and are with appropriate immune cell type. n = 12 (non-pregnant), n= 5 (1T), n=1 (2T), n= 3 (3T). uNK, uterine Natural Killer (**B)** Immune cells from each of the four stages in the reproductive cycle subset to 2200 cells. Immune cells separated by stage and then visualised by UMAP. Colours are indicative of clusters. **(C)** Using the scRNA seq dataset, graphs show frequency of total NK from CD45+ lymphocytes and then frequency of each uNK subset (uNK1, −2 −3) both as a proportion of CD45+ lymphocytes and proportion of total uNK cells. Means and standard deviations are shown for n = 3 (proliferative), n = 3 (secretory) n= 5 (1T), n=1 (2T), n= 3 (3T DB), n =3 (3T DP) DB, decidua basalis; DP, decidua parietalis; 1T, first trimester; 2T, second trimester; 3T, third trimester.

In the first trimester, uNK can be distinguished from circulating NK cells by their expression of CD49a and CD9; the subsets are then defined by their expression of CD39 and CD103 (22). We confirmed the presence of CD49a+ uNK in endometrium and in first and third trimester decidua, and that the three subpopulations uNK1, −2 and −3 can be identified using CD39 and CD103 (Fig 2a). However, in third trimester samples there was a significant CD49a+CD9-population. A comparison of CD49a+CD9+ and CD49a+ CD9-detected no phenotypic differences between these two populations, suggesting that CD49a alone can be used to identify uNK cells in the third trimester (Supp Fig 2). For consistency of gating strategy, we also identified uNK cells by their expression of CD49a alone in endometrial, first trimester and postpartum samples.

**Figure 2.**
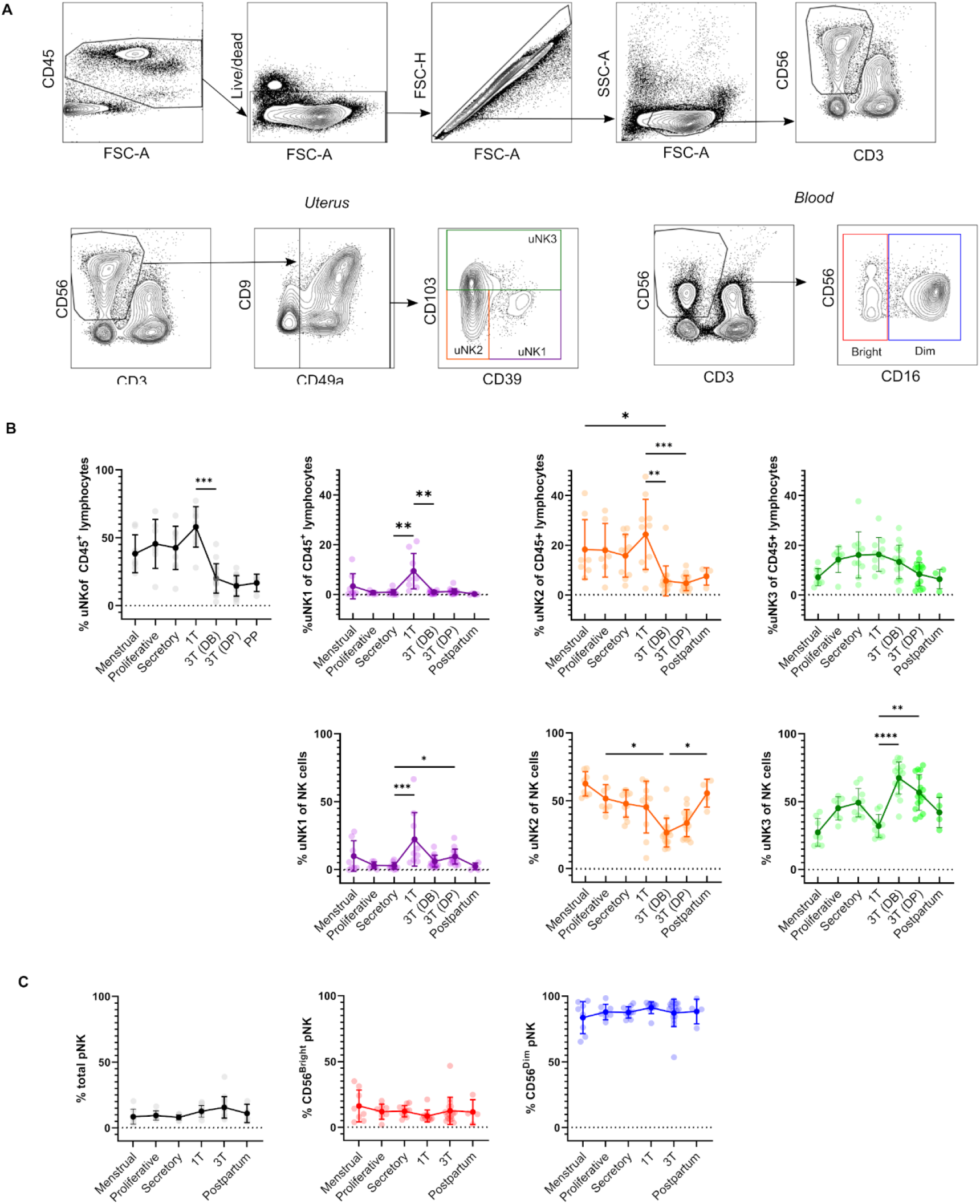
uNK1, −2 and −3 are present throughout the human reproductive cycle, by flow cytometry. **(A)** FACs gating strategy used to identify three uNK subsets and pNK (representative example shown). Coloured boxes in final plot indicate colour used for that subset in subsequent graphs. **(B)** Using flow cytometry data, graphs show frequency of total NK from CD45+ lymphocytes and then frequency of each uNK subset (uNK1, −2 −3) both as a proportion of CD45+ lymphocytes and proportion of total uNK cells. Means and standard deviations are shown for n = 8 (menstrual), n = 7 (proliferative), n = 10 (secretory) n= 10 (1T), n= 16 (3T DB), n = 16 (3T DP), n = 4 (postpartum). Statistical testing was done using Kruskal Wallis with a post-hoc Dunn test * p < 0.05, ** p < 0.01, *** p < 0.001 **** p< 0.0001 **(C)** Using flow cytometry data, graphs show frequency of total NK from CD45+ lymphocytes and then frequency of each pNK subset (CD56Brightand CD56Dim) as a proportion of total pNK. n numbers for each group are the same as **B.** DB, decidua basalis; DP, decidua parietalis; 1T, first trimester; 2T, second trimester; 3T, third trimester.

In line with previous reports (7), we observed a peak in total uNK, as a proportion of total CD45+ lymphocytes, in first trimester pregnancy by both scRNAseq and flow cytometry (Figure 1c, Fig 2b). We observed a similar proportion of total uNK in proliferative and secretory phase (Fig. 1c, 2c), in contrast to a previous report of higher proportion of uNK in secretory phase (38). This may be due to our use of CD49a allowing the removal of contaminating pNK from analysis, or due to their more vigorous sub-classification of the secretory stage.

Next, we examined uNK 1, −2 and −3 frequency expressed as either a proportion of total CD45+ lymphocytes or total uNK. We observed an increase in frequency of uNK1 when transitioning from secretory phase to first trimester pregnancy, but this was not sustained into third trimester decidua. This observation applied to both expression of uNK1 as percentage of both CD45+ lymphocytes and percentage of total NK cells. (Fig. 1c and Fig 2b), and the change was significant when measured by flow cytometry.

The variation of uNK2 frequency was similar to that observed for uNK1, with a peak in the first trimester observed by scRNAseq, and flow cytometry when frequency was measured as a percentage of CD45+ lymphocytes (Fig 1c and 2b). For the latter, uNK2 were significantly higher in the first trimester, compared to the third. Further, there was an upward trend of uNK2 when transitioning from third trimester decidua to postpartum endometrium when measured as a proportion of total NK (Fig 2c).

For uNK3, there was no change in frequency through the menstrual cycle. When measured as a percentage of CD45+ lymphocytes, there was a reduction in uNK3 in third trimester decidua parietalis, compared to both first trimester decidua and third trimester decidua basalis. This was significant when measured by flow cytometry. When measuring uNK3 as a percentage of total uNK there was a dip in the first trimester and a peak in both types of third trimester decidua. This was significant when measured by flow cytometry. The discrepancy between the proportions when expressed as a percentage of CD45+ lymphocytes or total uNK cells is likely due to the change in frequency of total uNK, as a proportion of CD45+ lymphocytes.

Within the third trimester decidua, the uNK2 population appeared greater in proportion of total uNK in the decidua parietalis compared to the decidua basalis in both scRNAseq and flow cytometry, although this did not reach significance for either (Fig 1c and 2b). The uNK3 population appeared greater in the decidua basalis, compared to decidua parietalis, which was significant when measured by flow cytometry as a percentage of CD45+ lymphocytes (Fig. 1c and 2b).

### Peripheral blood NK cell frequency does not vary over the reproductive cycle or correlate with uNK frequency

We also examined CD56Bright and Dim NK cells in matched peripheral blood by a conventional gating strategy to identify these populations. (Fig 2c). Unlike uNK, there was no variation in total CD56+pNK, CD56Bright or CD56Dim in peripheral blood when transitioning through different phases of the reproductive cycle. Furthermore, there was no significant correlation in levels of pNK and uNK subsets when expressed either as proportion of CD45+ live lymphocytes or total NK cells. (Supp Fig 3).

### uNK subsets upregulate KIR and LILRB1 during transition from non-pregnant endometrium to first trimester decidua

We next examined uNK expression of receptors that interact with trophoblast cells: KIR2DL1 and KIR2DL2/3 recognise HLA-C, LILRB1 recognises HLA-G and CD94 recognises HLA-E (39). In line with earlier findings on first trimester uNK (22, 24), we observed that uNK1 expressed higher levels of KIR than uNK2 and −3 (Fig 3b,c). We also found that all three uNK subsets expressed increased KIR in the first trimester of pregnancy, compared to non-pregnant endometrium and third trimester decidua (Fig 3b,c). Similar to KIR, LILRB1 protein expression peaked in the first trimester, although this was only statistically significant in uNK2 and uNK3 (Fig 3b). *LILRB1* transcript expression followed a similar trend, although in contrast to our findings at the protein level, *LILRB1* mRNA was not detectable in the decidua basalis (Fig 3c). At the transcript and protein level, CD94 was expressed at a higher level on uNK2 and 3 compared to uNK1 (Fig.3b, c). There was a slight reduction in CD94 transcript (*KLRD1*) towards the end of pregnancy, but this was not observed at the protein level. (Fig. 3b, c)

**Figure 3.**
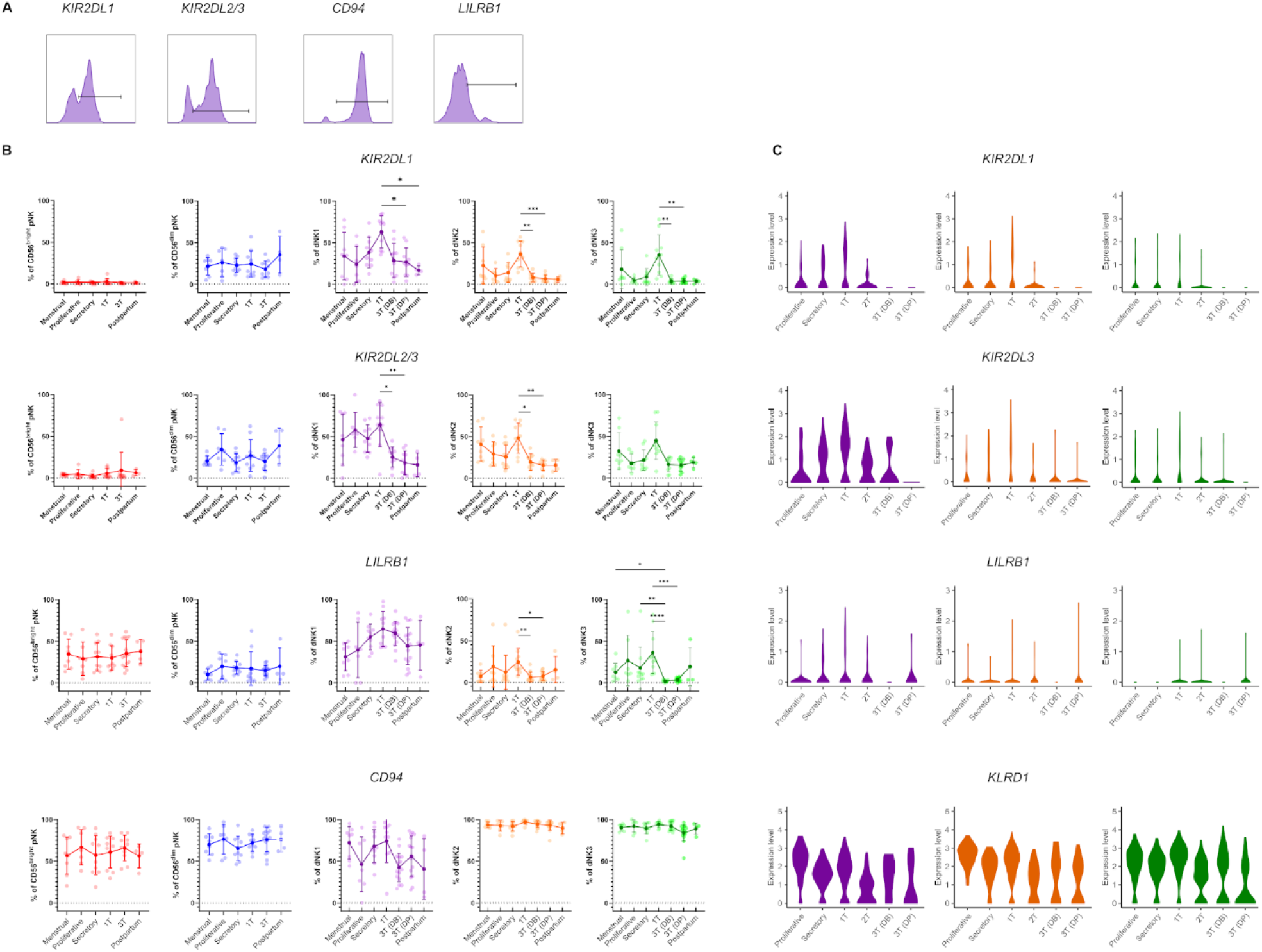
uNK upregulate expression of KIR and LILRB1 in first trimester. **(A)** Uterine and peripheral NK cells taken at different stages of the reproductive cycle were freshly stained for phenotypic markers. Representative staining from secretory phase uNK1 is shown. The positive gates were set by reference to FMO controls. **(B)** Graphs showing frequencies of KIR2DL1, KIR2DL2/3, LILRB1 and CD94 on pNK and uNK subsets. Means and standard deviations are shown for n = 8 (menstrual), n = 7 (proliferative), n = 10 (secretory) n= 10 (1T), n= 16 (3T DB), n = 16 (3T DP), n = 4 (postpartum). Statistical testing was done using Kruskal Wallis with a post-hoc Dunn test * p < 0.05, ** p < 0.01, *** p < 0.001. **(C)** Violin plots showing corresponding mRNA expression in uNK subsets over the reproductive cycle as determined by scRNAseq. DB, decidua basalis; DP, decidua parietalis; 1T, first trimester; 2T, second trimester; 3T, third trimester.

In line with our finding that pNK did not change in frequency over the reproductive cycle, examination of NK cells from matched blood showed no change in the frequency at which KIR, LILRB1, and CD94 are expressed in these cells. (Fig. 3b).

### uNK are the most active at the time of implantation

We next assessed functional responses with and without stimulation with PMA and ionomycin (Fig. 4a). CD107a staining is a proxy for degranulation and previous studies have shown that this acts as a reliable measure of overall uNK activation (40). We also examined the production of IL-8 and GM-CSF, which are thought to promote EVT invasion (6, 40, 41), and the classical NK cell cytokines IFNγ and TNFα (Fig. 4b).

**Figure 4.**
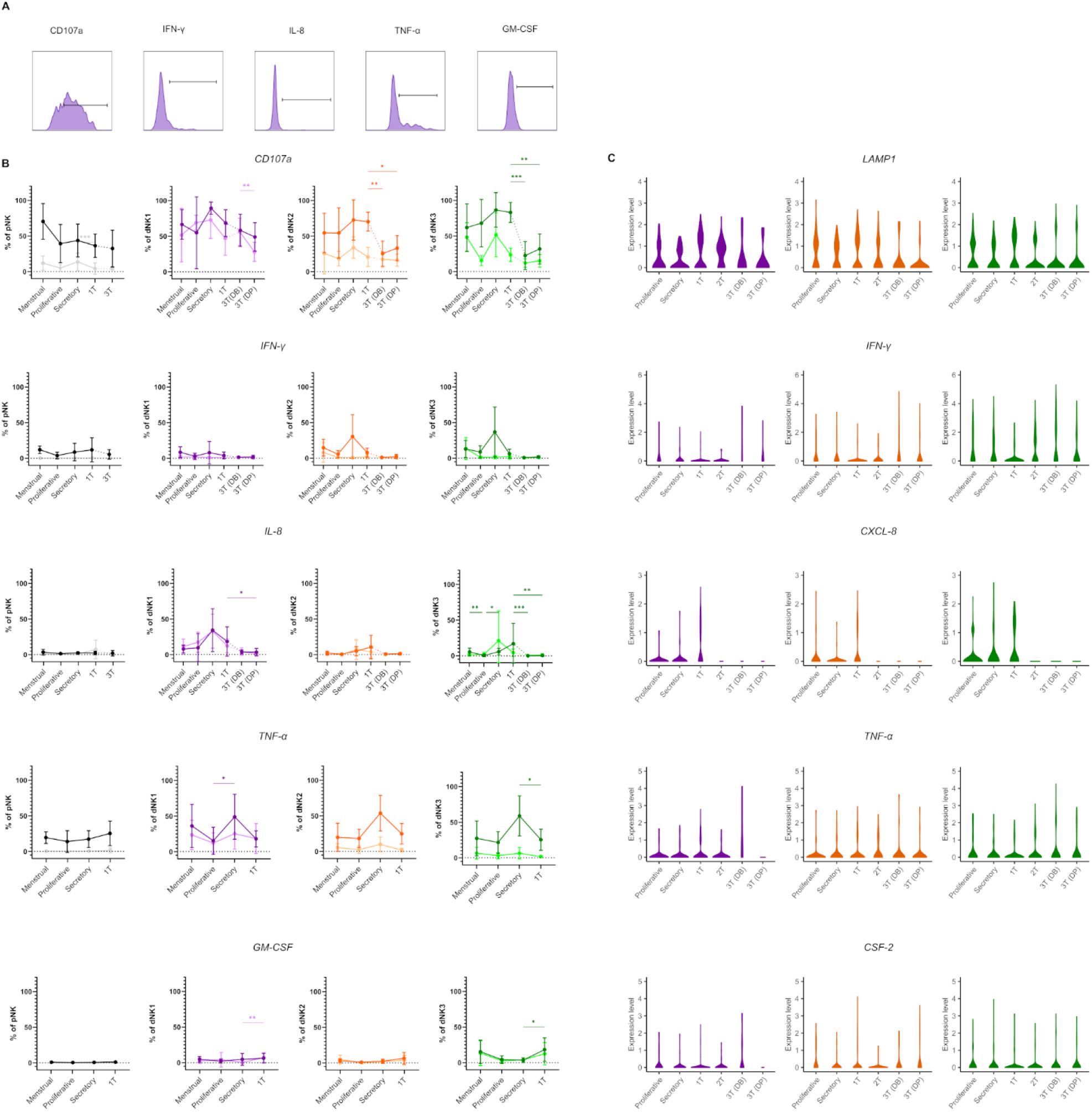
uNK are most active at the time of implantation. **(A)** Uterine and peripheral NK cells taken at different stages of the reproductive cycle were cultured with or without PMA and ionomycin stimulation and assessed for degranulation (CD107a) and production of IFNγ, IL-8, TNFα and GM-CSF. Representative staining from secretory phase uNK1 is shown. The positive gates were set by reference to FMO controls. **(B)** Graphs showing frequencies of CD107a, IFNγ, IL-8, TNFα and GM-CSF on pNK and uNK subsets. Unstimulated cells are represented by lighter lines and stimulated cells by darker lines. Means and standard deviations are shown for n = 5 (menstrual), n = 6 (proliferative), n = 10 (secretory) n= 10 (1T), n= 16 (3T DB), n = 16 (3T DP). Statistical testing was done using Kruskal Wallis with a post-hoc Dunn test * p < 0.05, ** p < 0.001, *** p < 0.0001. **(C)** Violin plots showing corresponding mRNA expression in uNK subsets over the reproductive cycle as determined by scRNAseq. DB, decidua basalis; DP, decidua parietalis; 1T, first trimester; 2T, second trimester; 3T, third trimester.

Degranulation in unstimulated conditions declined during the proliferative phase, slightly in uNK2 and significantly in uNK3, compared the other two phases of the menstrual cycle. (Fig. 4b). At the end of pregnancy, degranulation was significantly lower in third trimester decidua parietalis compared to decidua basalis in dNK1, but this was not replicated in the other subsets. (Fig. 4b). In stimulated cells there was a reduction in degranulation in uNK2 and uNK3 in both third trimester decidua compared to first trimester decidua (Fig. 4b).

For TNF-α, IFN-γ and IL-8, we observed peaks in cytokine production across all stimulated uNK subsets during secretory phase, compared to the proliferative phase and first trimester, although this did not reach statistical significance in all cases (Fig 4b). For IL-8 in uNK3 this peak was maintained into first trimester pregnancy. This trend was also present in unstimulated cells for IL-8 (Fig 4b).

Third trimester uNK produced less cytokine than first trimester uNK, although this only reached significance in IL-8 production from uNK3. This reduction in cytokine production through pregnancy was also seen at the mRNA level for IL-8 (Fig. 4c). In contrast, GM-CSF protein production was consistently low in the menstrual cycle, including the secretory phase, but increased significantly in the first trimester of pregnancy in unstimulated uNK1 and stimulated uNK3. (Fig. 4b).

For examination of NK cells from matched peripheral blood, data from CD56Brights and Dims are shown together due to downregulation of CD16 after stimulation. Aside from a significant decline in CD107a expression in unstimulated cells when transitioning from secretory phase to first trimester pregnancy, there were no distinct trend in both stimulated and unstimulated cells (Fig. 4b). The mRNA expression of other NK cell proteins of interest across the reproductive cycle, such as granzymes, that were not included in the flow cytometry panel, can be seen in the supplementary figures 4 and 5.

### uNK phenotype and function do not change in labour

In line with previous findings (42, 43), we did not observe any change in the proportion of total uNK in labouring compared to non-labouring decidua (Fig 5b). The proportion of uNK in non-labouring decidua was lower in the decidua parietalis compared to the decidua basalis (Fig 5b). This is in contrast to a previous report which found a higher frequency of CD56 bright NK cells in the parietalis than the basalis (44). However, the lack of tissue-specific markers means that it is difficult to be sure if these all represent uNK.

**Figure 5.**
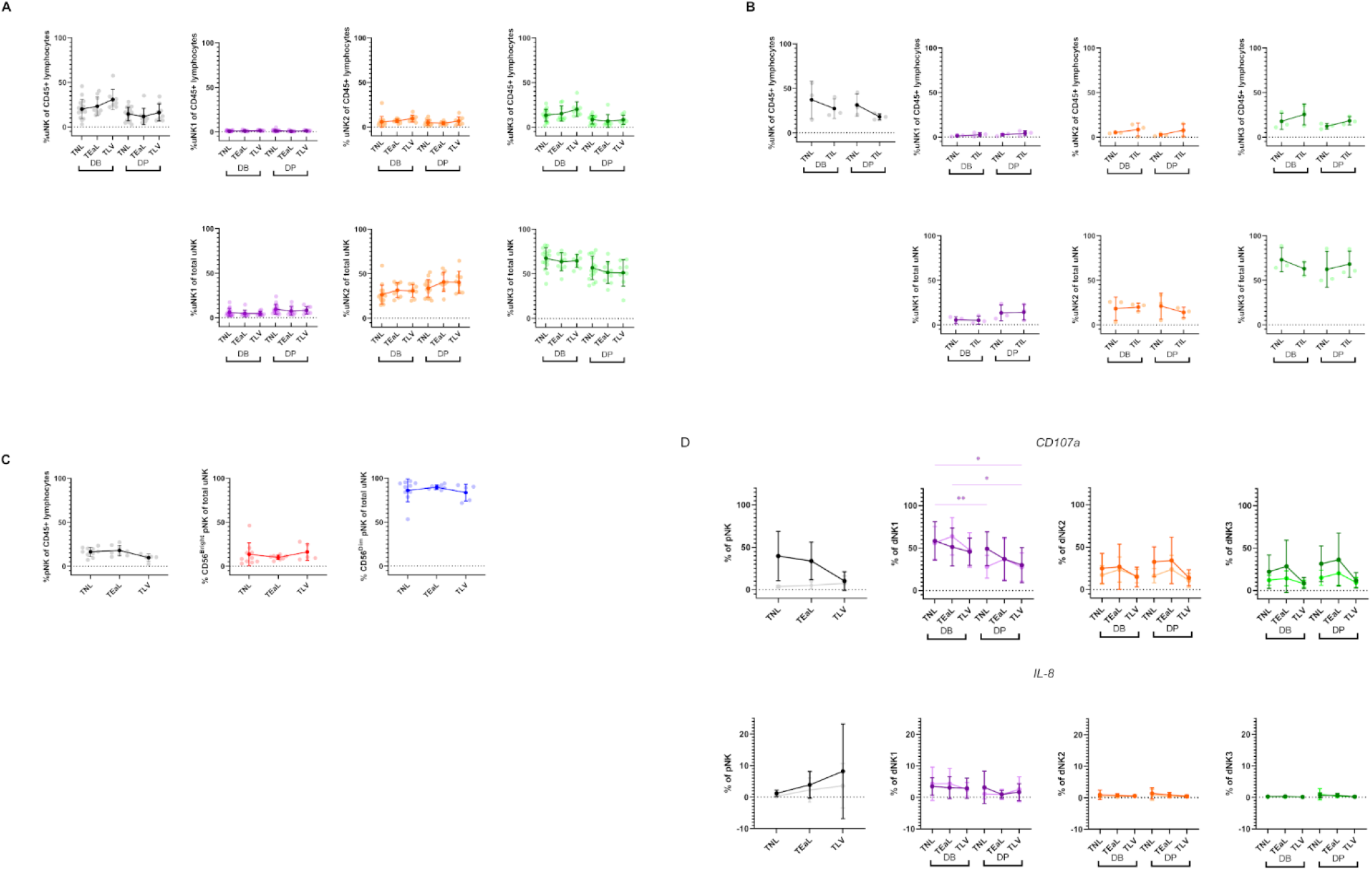
uNK phenotype and function do not change in labour. **(A)** The FACs gating strategy shown in **Figure 2A** was used to identify the three uNK subsets. Graphs show frequency of total NK from CD45+ lymphocytes and then frequency of each uNK subset (uNK1, −2 −3) both as a proportion of CD45+ lymphocytes and proportion of total uNK cells. Graphs are divided to show results from both decidua basalis and decidua parietalis. Means and standard deviations are shown for n = 16 (TNL), n = 9 (TEaL), n = 9 (TLV). Statistical testing was done using Kruskal Wallis with a post-hoc Dunn test * p < 0.05. (**B**) Using the scRNA seq dataset, graphs show frequency of total NK from CD45+ lymphocytes and then frequency of each uNK subset (uNK1, −2 −3) both as a proportion of CD45+ lymphocytes and proportion of total uNK cells. Graphs are divided to show results from both decidua basalis and decidua parietalis. Means and standard deviations are shown for n = 3 (TNL), n = 3 (TIL). (**C**) The FACs gating strategy shown in Figure 2C was used to identify the two pNK subsets. Using flow cytometry data, graphs show frequency of total NK from CD45+ lymphocytes and then frequency of each pNK subset (CD56Bright and CD56Dim) from total NK. n numbers for each group are the same as **(A)**. **(D)** Uterine and peripheral NK cells were cultured as described in **Figure 4A**. Graphs showing frequencies of CD107a and IL-8 on pNK and uNK subsets. Unstimulated cells are represented by lighter lines and stimulated cells by darker lines. n numbers for each group are the same as **(A).** Statistical testing was done using Kruskal Wallis with a post-hoc Dunn test * p < 0.05. DB, decidua basalis; DP, decidua parietalis; TNL, term non-labouring; TEaL, term early labouring; TLV, term vaginal birth; TIL, term in labour

We next examined the uNK subpopulations in non-labour compared to early labour and established labour samples. We observed no change in the frequency of any of the uNK or pNK subsets across the spectrum of these samples (Fig. 5a and 5b). The receptors examined were also stable during labour (Supp Fig 6), an observation that suggests that EVT cross-talk with uNK is not a major participant in labour. Similarly, for those markers examined, the function of uNK and pNK subsets remain stable during labour, although we did observe that, regardless of labouring state, the uNK1 population in the decidua basalis is significantly more active than the population in the decidua parietalis (Figure 5 and supp fig 7). This is supported by the low number of differentially expressed genes across all three uNK subsets in the scRNAseq data comparing non-labour to labour samples (Supp table 3).

## Discussion

To our knowledge, this is the first study to track the three uNK subpopulations throughout the reproductive cycle, looking at their frequency, phenotypes and functions. In line with previous studies, we found that the total uNK population peaked during the first trimester of pregnancy (7, 45). We discovered that uNK1 and uNK2 peak in this period, but that uNK3 peaks towards the end of pregnancy. This aligns with a recent report that KIR+CD39+ uNK (mostly representing uNK1) and KIR+CD39-(mostly representing uNK2) increase in frequency towards the end of the menstrual cycle and remain elevated in early pregnancy (46). Both these reports support the proposal that uNK1 communicate with EVTs in early pregnancy (22, 24), but may also point to a role for uNK2 in this process.

It has previously been reported that KIR expression by total uNK increases in the first trimester, compared to non-pregnant endometrium (21). Our finding that the major KIR-expressing subset, uNK1, is most frequent in the first trimester might suggest that this increase in KIR expression is at least partially a result of greater prominence of uNK1 but interestingly, all three uNK subsets express increased KIR in the first trimester of pregnancy. A similar trend was seen for LILRB1. This suggests that all uNK subpopulations increase their ability to recognise EVT, via HLA-C and HLA-G, in the first trimester. In comparison, CD94 expression was higher on uNK2 and −3 subsets compared to uNK1. This is in contrast with previous findings (24). However, this measured marker intensity on recovered cryopreserved cells, whereas we report percentage of CD94+ fresh cells. Therefore, it is possible that the freezing process preferentially killed CD94-uNK1 cells or uNK1 have a lower percentage of cells expressing CD94, but have a higher expression per cell.

By examining degranulation as a proxy for general NK cell activation (40), we found that uNK were typically most active at around the time of implantation, in the secretory phase of the menstrual cycle and the first trimester of pregnancy. This is in line with previous findings that first trimester uNK are more able to degranulate in response to HCMV-infected targets than those at term, although the same study found that, following IL-15 stimulation, term uNK are better able to degranulate in response to PMA and ionomycin than first trimester cells (9). Without stimulation, uNK produce little IFNγ, IL-8, TNFα and GM-CSF. However, after stimulation with PMA and ionomycin, uNK have the highest ability to produce most of these cytokines during the secretory phase, except for GM-CSF, whose production peaks in the first trimester. This finding is interesting because the timing of maximum activation coincides with the window of implantation, suggesting that uNK may have a role in coordinating successful implantation.

Cumulative evidence indicates an important physiological role for uNK in first trimester pregnancy, but there is conflicting evidence on their role in reproductive failure. Our findings and those of others (22, 24) point to uNK1 as the uNK subset more likely to mediate placental implantation in early pregnancy. Future studies focusing specifically on this subset may be able to elucidate differences that were previously masked due to examination of uNK as a bulk population. A recent study using scRNAseq suggests that there is a reduction of uNK1 in pathological pregnancies (47); however, these findings need to be interpreted with caution because the pathological samples were collected after pregnancy loss, making it difficult to discern if changes seen in immune cells are a cause or an effect, due to inflammatory changes that typically occur after fetal demise. To overcome this, future studies could interrogate uNK during window of implantation (48) or from elective termination of pregnancy samples stratified to low and high risk by uterine artery doppler which has high specificity in predicting risk of pre-eclampsia and intrauterine growth restriction (49).

In the third trimester the majority of uNK cells are uNK3, which express low levels of KIRs and LILRB1. This could suggest that, in contrast to early pregnancy, the major role for uNK in late pregnancy does not involve interactions with EVTs. Similarly, the expression of the functional markers we examined was lower in the third trimester. This could indicate that, if these cells have a role at the end of pregnancy, it is via a different mechanism of action. Intriguingly, by scRNAseq, uNK3 were the most transcriptionally different between non-labouring and labouring states, suggesting they may have a role in labour that is yet to be defined. It would be interesting to do a broader analysis of these cells, or examine how they look in pathological cases such as pre-eclampsia or preterm birth.

In conclusion, we show here how uNK subset number, expression of receptors and function change dynamically across the healthy reproductive cycle. This provides evidence on their physiological role in implantation, but will also provide an important platform from which the relationship between uNK function and pathologies of pregnancy associated with impaired implantation and placentation can be investigated.

## Availability of data and code

For scRNAseq analyses, data is available from: http://www.reproductivecellatlas.org/ (non-pregnant uterus), Array Express E-MTAB-6701 (first trimester), dbGaP phs001886.v1.p1 (second and third trimester). Code is available from https://github.com/ewhettlock/reproductive_cycle. Flow cytometry data is available from https://osf.io/wkxyz/.

## Author Contributions

EMW, EVW and VM designed the study, analysed results and wrote the manuscript. EMW, EVW and AOC carried out experiments. EVW, BB and MRJ consented patients and collected clinical samples. All others contributed to editing the manuscript.

## Acknowledgements

We would like to thank all the people from Chelsea and Westminster Hospital, West Middlesex University Hospital and John Hunter Clinic (London, UK) who contributed samples to this study. We would also like to thank Dr Pei Lai, Dr Nishel Shah, Miss Sharmista Guha and all the clinical staff who helped in the collection of samples.

