## Supplementary information for "Dynamic changes in uterine NK cell subset frequency and function over the menstrual cycle and pregnancy" (from another version of this preprint: version 1, 2022-02-21)

### *Whettlock, Woon et al - Supplementary Material*

**Supplementary Table 1.** Demographics from non-pregnant participants

| Patient | Age | Parity | Cycle day | Cycle length | Serum progesterone (ng/mL) | Phase assigned | Assessment |
|---|---|---|---|---|---|---|---|
| 1 | 38 | Multiparous | 1 | 28 | <1 | Menstrual | Phenotype |
| 2 | 36 | Multiparous | 1 | 28 | 3 | Menstrual | Phenotype and function |
| 3 | 23 | Primiparous | 2 | 28 | 2 | Menstrual | Phenotype and function |
| 4 | 25 | Primiparous | 1 | 28 | <1 | Menstrual | Phenotype |
| 5 | 21 | Primiparous | 5 | 28 | 1 | Menstrual | Phenotype and function |
| 6 | 26 | Primiparous | 2 | 24 | Not taken | Menstrual | Phenotype and function |
| 7 | 38 | Multiparous | 1 | 32 | 1 | Menstrual | Phenotype and function |
| 9 | 24 | Primiparous | 2 | 30 | 1 | Menstrual | Phenotype |
| 10 | 29 | Primiparous | 10 | 35 | 6 | Proliferative | Phenotype |
| 11 | 29 | Primiparous | 7 | 28 | <1 | Proliferative | Phenotype and function |
| 12 | 35 | Multiparous | 8 | 30 | 4 | Proliferative | Phenotype |
| 13 | 21 | Primiparous | 10 | 28 | Not taken | Proliferative | Phenotype and function |
| 14 | 31 | Multiparous | 10 | 28 | 5 | Proliferative | Phenotype and function |
| 15 | 25 | Primiparous | 11 | 28 | <1 | Proliferative | Phenotype and function |

| | | | | | | | |
|---|---|---|---|---|---|---|---|
| 16 | 34 | Primiparous | 13 | 28 | Not taken | Proliferative | Phenotype and function |
| 17 | 27 | Primiparous | 21 | 30 | 30 | Secretory | Phenotype |
| 18 | 28 | Primiparous | 20 | 21 | 2 | Secretory | Phenotype |
| 19 | 27 | Primiparous | 17 | 27 | 11 | Secretory | Phenotype and function |
| 20 | 20 | Primiparous | 15 | 28 | 7 | Secretory | Phenotype and function |
| 21 | 30 | Primiparous | 19 | 28 | 51 | Secretory | Phenotype and function |
| 22 | 25 | Primiparous | 17 | 28 | Not taken | Secretory | Phenotype and function |
| 23 | 33 | Multiparous | 19 | 30 | 22 | Secretory | Phenotype and function |
| 24 | 40 | Multiparous | 21 | 28 | 28 | Secretory | Phenotype and function |
| 25 | 32 | Primiparous | 17 | 28 | 13 | Secretory | Phenotype and function |
| 26 | 27 | Primiparous | 20 | 28 | 23 | Secretory | Phenotype and function |
| 27 | 27 | Primiparous | 20 | 42 | 7 | Secretory | Function |
| 28 | 44 | Multiparous | N/A | N/A | <1 | Postpartum 16 weeks | Phenotype |
| 29 | 35 | Multiparous | N/A | N/A | <1 | Postpartum 10 weeks | Phenotype |
| 30 | 37 | Multiparous | N/A | N/A | 3 | Postpartum 4 weeks | Phenotype |
| 31 | 29 | Multiparous | N/A | N/A | <1 | Postpartum 6 weeks | Phenotype |

**Supplementary Table 2.** Demographics from pregnant participants (first and third trimester)

| Patient | Age | Self-reported Ethnicity | Gravida and Parity | Gestation | Phase | Reason for Caesarean Section | Assessment |
|---|---|---|---|---|---|---|---|
| 1 | 17 | Asian | G2P1 | 11+2 | 1T | N/A | Phenotype only |
| 2 | 24 | White | G2P1 | 8+ 6 | 1T | N/A | Phenotype and function |
| 3 | 40 | White | G6P4 | 9+1 | 1T | N/A | Phenotype and function |
| 4 | 20 | White | G1P0 | 6+6 | 1T | N/A | Phenotype and function |
| 5 | 32 | White | G3P0 | 10+4 | 1T | N/A | Phenotype and function |
| 6 | 25 | Asian | G1P0 | 11+5 | 1T | N/A | Phenotype and function |
| 7 | 22 | White | G1P0 | 8+3 | 1T | N/A | Phenotype and function |
| 8 | 21 | White | G3P0 | 13+0 | 1T | N/A | Phenotype and function |
| 9 | 33 | White | G3P1 | 12+6 | 1T | N/A | Phenotype and function |
| 10 | 40 | Asian | G5P4 | 8+6 | 1T | N/A | Phenotype and function |
| 11 | 40 | White | G4P3 | 10+4 | 1T | N/A | Function only |
| 12 | 33 | White - Canadian | G2P1 | 39+0 | TNL | Previous CS | Phenotype and function |
| 13 | 39 | Middle Eastern - Turkish | G3P2 | 38 + 0 | TNL | Previous CS | Phenotype and function |
| 14 | 36 | White British | G5P1 | 37 + 0 | TNL | Maternal request | Phenotype and function |
| 15 | 39 | White - Italian | G4P2 | 38+6 | TNL | Previous CS | Phenotype and function |
| 16 | 37 | White - French | G2P1 | 38 + 0 | TNL | Previous CS | Phenotype and function |
| 17 | 35 | White - British | G4P1 | 39 + 0 | TNL | Previous CS | Phenotype and function |
| 18 | 37 | White - Polish | G3P1 | 39+1 | TNL | Previous CS | Phenotype and function |
| 19 | 33 | White British | G1P0 | 39+0 | TNL | Fetal medical condition | Phenotype and function |
| 20 | 40 | White – Eastern European | G2P1 | 39+3 | TNL | Previous CS | Phenotype and function |
| 21 | 34 | White - Greek | G1P0 | 39+0 | TNL | Breech | Phenotype and function |
| 22 | 33 | White - British | G6P0 | 39+1 | TNL | Maternal request | Phenotype and function |
| 23 | 34 | East African - Ethiopian | G3P2 | 39+4 | TNL | Previous CS | Phenotype and function |
| 24 | 31 | White - British | G2P1 | 39+2 | TNL | Breech | Phenotype and function |
| 25 | 35 | White - British | G5P4 | 39+2 | TNL | Previous CS | Phenotype and function |
| 26 | 31 | White - British | G3P1 | 37+1 | TNL | Maternal request | Phenotype and function |
| 27 | 33 | Iranian | G2P1 | 39+1 | TNL | Previous CS | Phenotype and function |

| | | | | | | | |
|---|---|---|---|---|---|---|---|
| 28 | 31 | White – British | G6P1 | 39 + 3 | TEaL | Maternal request | Phenotype and function |
| 29 | 38 | Asian – Pakistani | G1P0 | 38+3 | TEaL | Maternal request | Phenotype and function |
| 30 | 35 | White – British | G2P1 | 39+2 | TEaL | Maternal request | Phenotype and function |
| 31 | 33 | White – United States | G2P0 | 41+1 | TEaL | Breech | Phenotype and function |
| 32 | 29 | Mixed race – White and South African | G2P0 | 40+5 | TEaL | Fetal distress | Phenotype and function |
| 33 | 34 | White – British | G1P0 | 38+3 | TEaL | Maternal request | Phenotype and function |
| 34 | 32 | White – United States | G1P0 | 38+5 | TEaL | Maternal request | Phenotype and function |
| 35 | 36 | Mixed race - Brazilian | G3P0 | 41+0 | TEaL | Previous CS | Phenotype and function |
| 36 | 35 | Asian - Indian | G3P0 | 39+2 | TEaL | Maternal request | Phenotype and function |
| 37 | 37 | White - British | G1P0 | 40+2 | TLV | N/A | Phenotype and function |
| 38 | 31 | White - Irish | G1P0 | 39+5 | TLV | N/A | Phenotype and function |
| 39 | 37 | Mixed Asian | G3P1 | 40+1 | TLV | N/A | Phenotype and function |
| 40 | 31 | South European – Italian | G1P0 | 40+6 | TLV | N/A | Phenotype and function |
| 41 | 29 | White - British | G1P0 | 39+6 | TLV | N/A | Phenotype and function |
| 42 | 31 | White - British | G1P0 | 41+5 | TLV | N/A | Phenotype and function |
| 43 | 34 | Caucasian - Czech | G2P1 | 40+4 | TLV | N/A | Phenotype and function |
| 44 | 36 | Asian – Chinese Malaysian | G3P1 | 40+3 | TLV | N/A | Phenotype and function |
| 45 | 36 | Lithuanian | G2P1 | 39+5 | TLV | N/A | Phenotype and function |

1T, first trimester; TNL term non labour; TEaL, term in early labour; TLV, term labour vaginal; CS = caesarean section

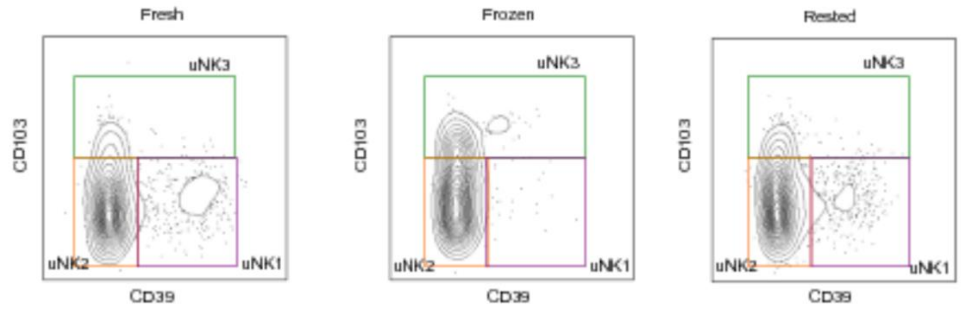


**Supplementary Figure 1.** Comparison of uNK from first trimester pregnancy, stained and acquired fresh, after cryopreservation and recovery, or after 16 hours rest. All treatments are from the same participant.

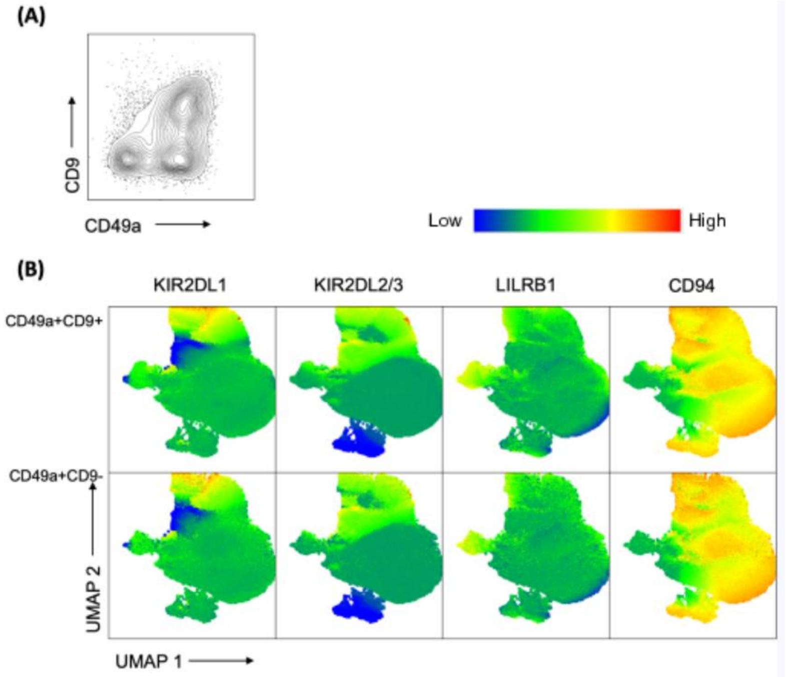


**Supplementary Figure 2.** (A) Representative staining of CD49a versus CD9 on CD56+CD3- live lymphocytes. (B) CD49a+CD56+CD3- live lymphocytes were concatenated from endometrial, first trimester and third trimester samples. They were gated as CD49a+ CD9+ and CD49a+ CD9- and the expression of the phenotypic markers (KIR23DL1, KIR2DL2/3, LILIRB1 and CD94) was visualized as a heatmap using UMAP plots.

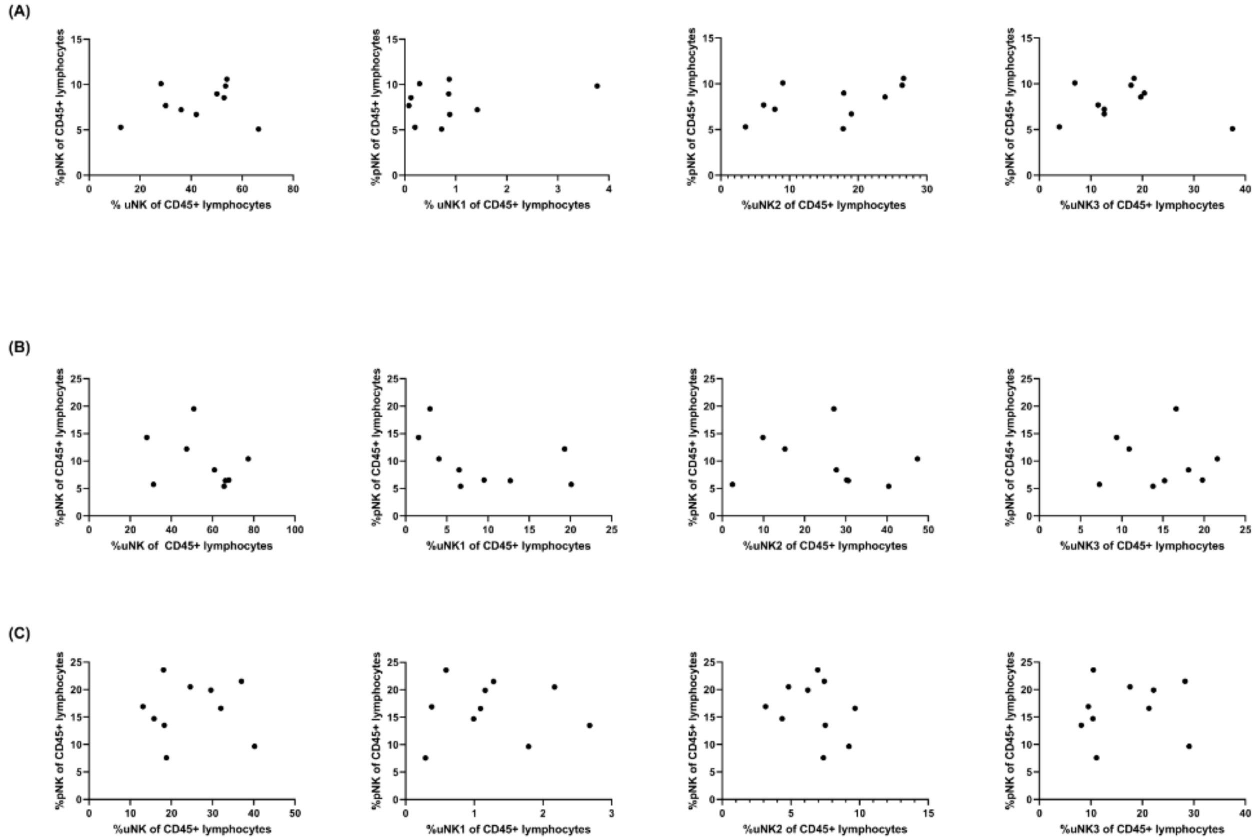


**Supplementary Figure 3.** Correlation between peripheral and uterine NK cells in secretory phase of the menstrual cycle (A), first trimester (B) and third trimester (C). No significant correlation was found between pNK and uNK, uNK1, uNK2 and uNK3. Additionally, no correlation between CD56$^{bright}$ and -$^{dim}$ and uNK1, uNK2 and uNK3 was found in all phases above when expressed as a percentage of CD45+ lymphocytes (not shown). Statistical testing was done by Spearman's rank correlation.

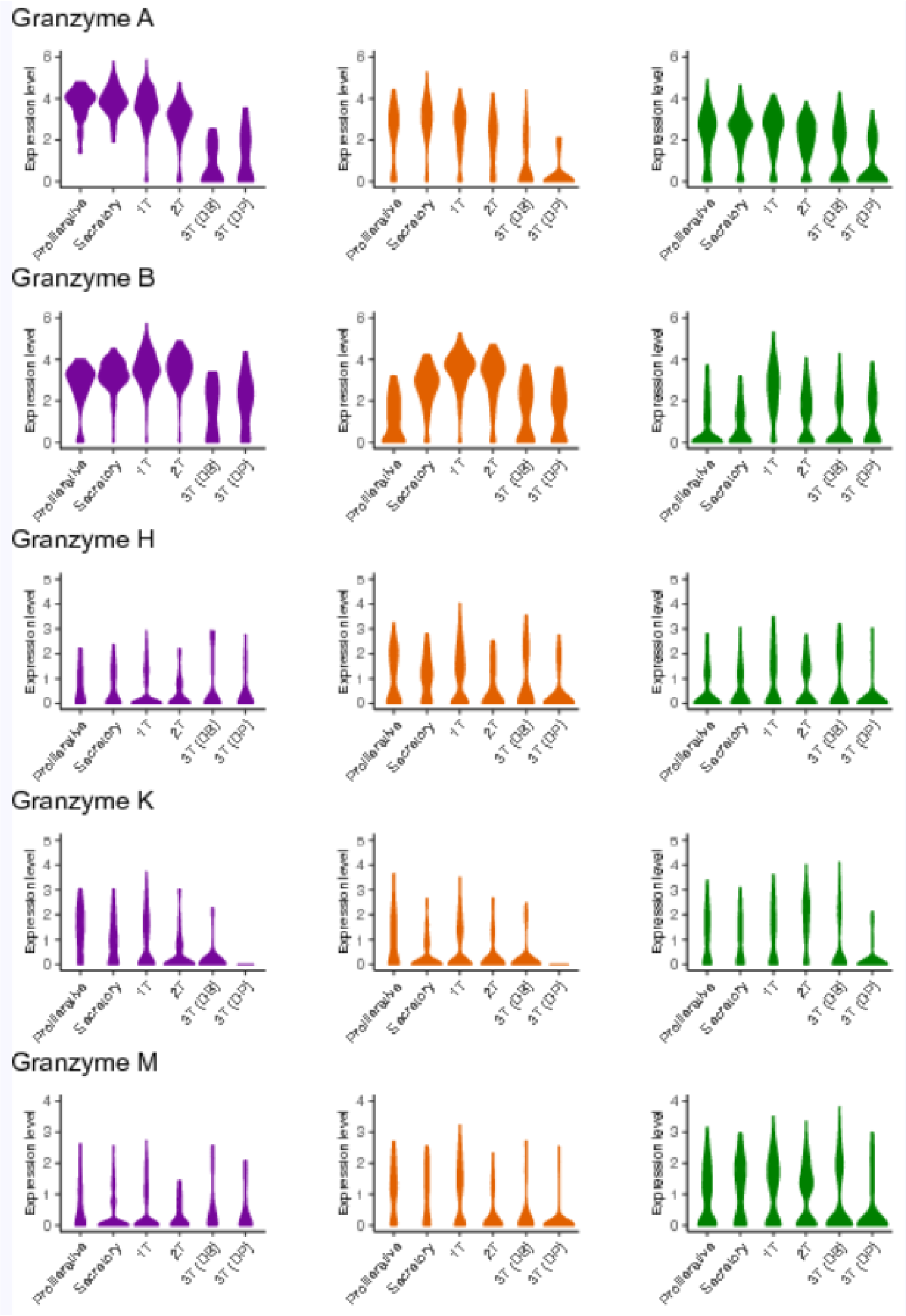


**Supplementary Figure 4.** Violin plots showing mRNA expression of the three uNK subsets (uNK1 in purple, uNK2 in orange, uNK3 in green) over six phases of the reproductive cycle.

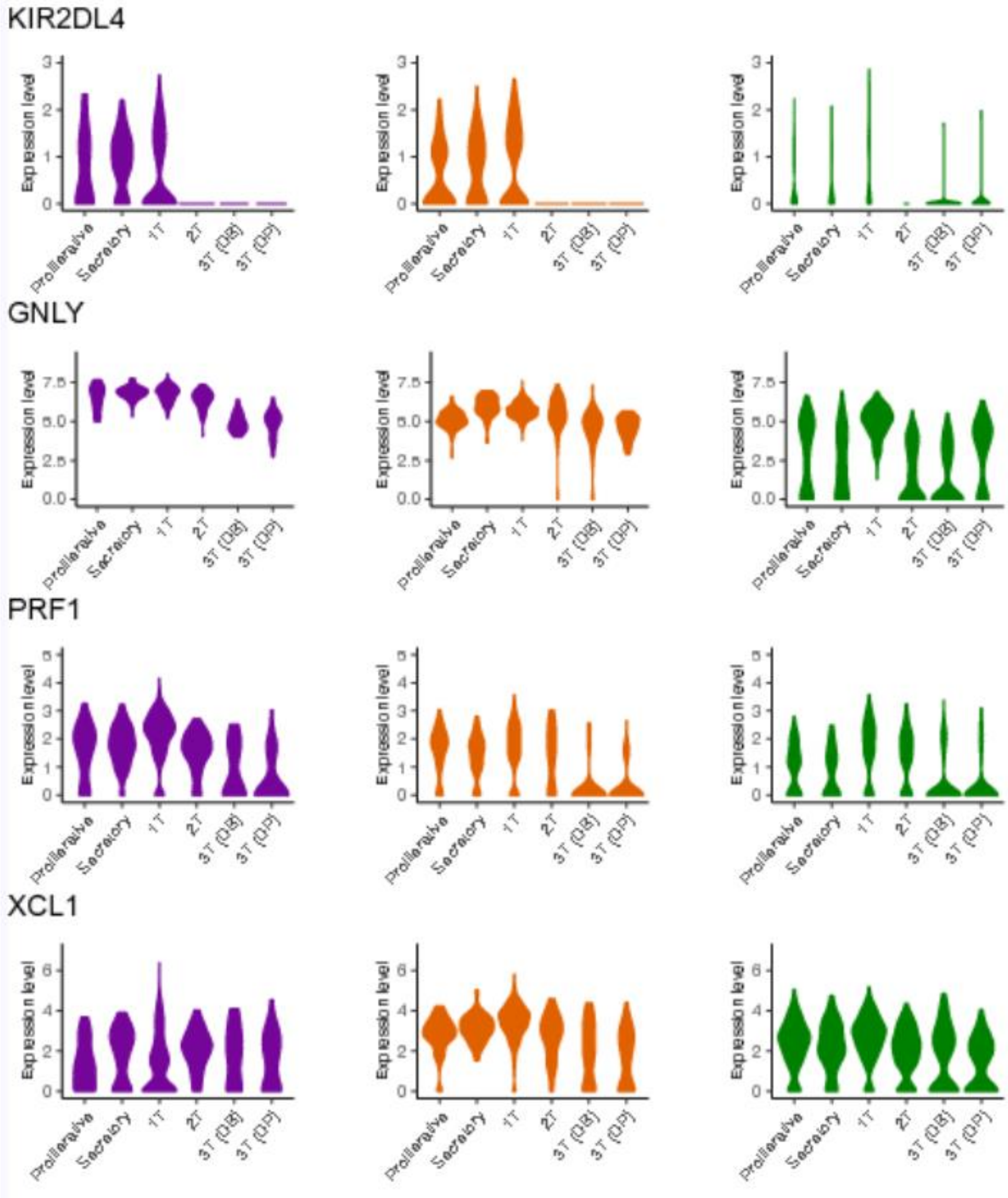


**Supplementary Figure 5.** Violin plots showing mRNA expression of additional markers of interest in the three uNK subsets (uNK1 in purple, uNK2 in orange, uNK3 in green) across six phases of the reproductive cycle.

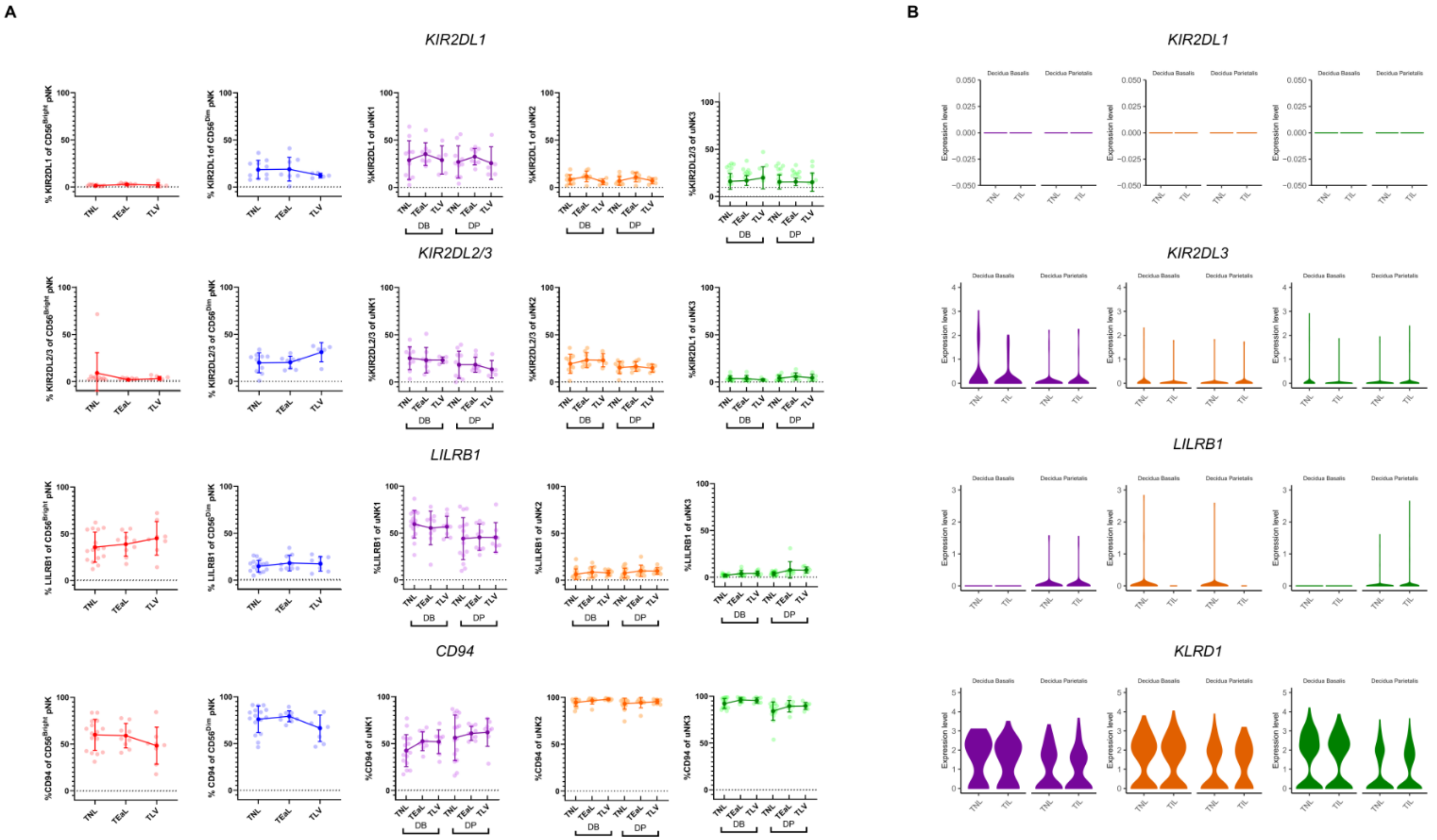


**Supplementary Figure 6.** Phenotype of NK cells in labour. (A) Graphs showing phenotypes of pNK and uNK in third trimester, acquired by flow cytometry. (B) Violin plots showing corresponding mRNA expression of receptors on uNK, acquired by scRNAseq. TNL, term not in labour; TEaL, term early labour; TLV, term in labour (vaginal); DB, decidua basalis; DP; decidua parietalis.

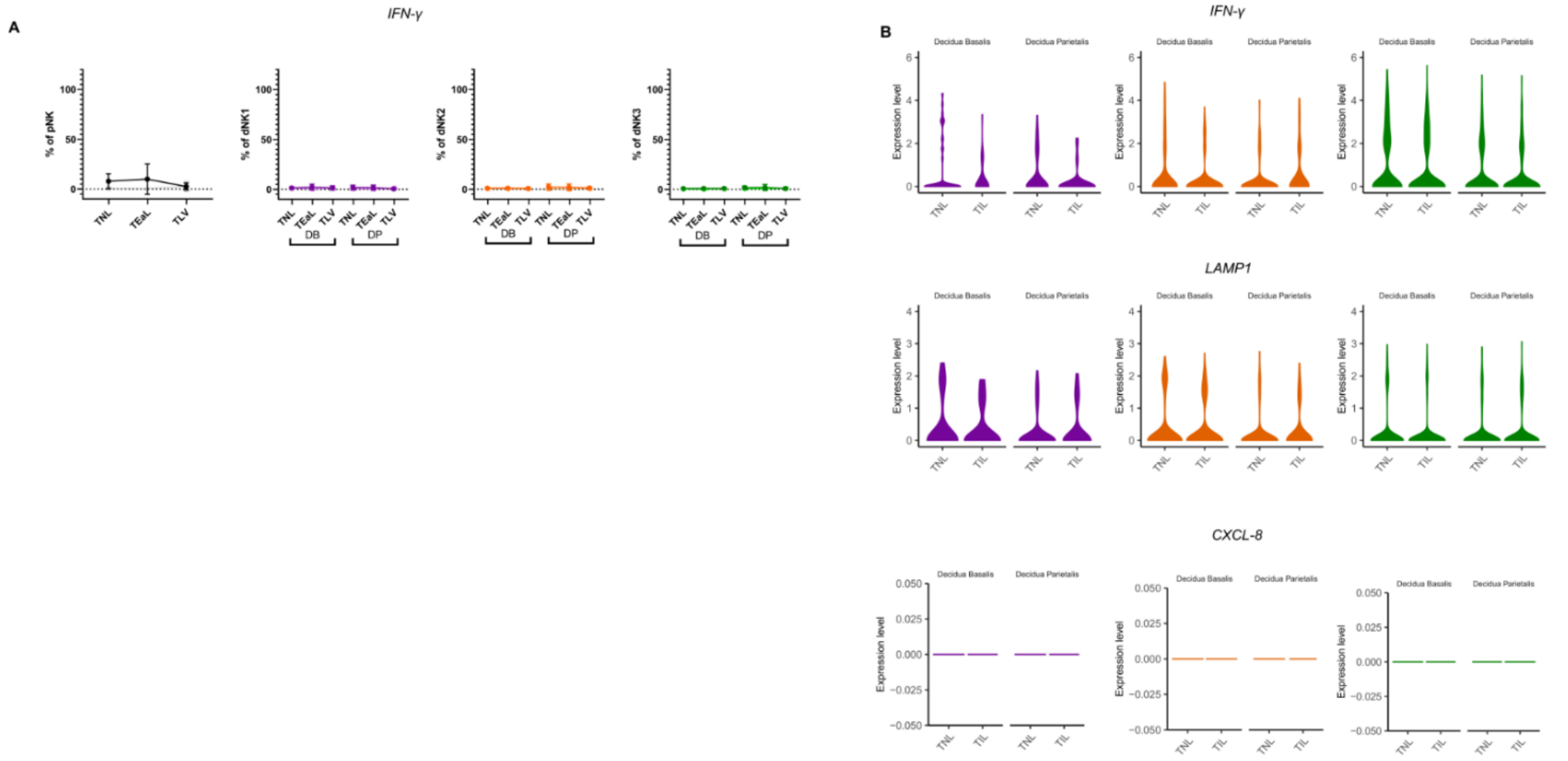


**Supplementary Figure 7.** Function of NK cells in labour. (A) Graphs showing frequencies of IFNγ in pNK and uNK acquired by flow cytometry in bulk uNK (black), uNK1 (purple), uNK2 (orange) and uNK3 (green). Unstimulated cells are represented by lighter lines and stimulated cells by darker lines. (B) Violin plots showing mRNA expression of IFNG, LAMP1 and CXCL8 in uNK1 (purple), uNK2 (orange) and uNK3 (green). TNL, term not in labour; TEaL, term early labour; TLV, term in labour (vaginal); TIL, term in labour.

**Supplementary Table 3** contains a large amount of data and is provided as a separate Excel file.
